# Deep Collection of Quantitative Nuclear Division Dynamics Data in RNAi-treated *Caenorhabditis elegans* Embryos

**DOI:** 10.1101/2020.10.04.325761

**Authors:** Koji Kyoda, Hatsumi Okada, Hiroya Itoga, Shuichi Onami

## Abstract

Recent advances in bioimage informatics techniques have yielded quantitative data on multicellular dynamics from microscopy images of animal development. Several such data collections have been created for *Caenorhabditis elegans* embryos under various gene silencing conditions. However, because of the limited depth of the datasets, it is impractical to apply standard statistical methods to these collections. Here, we created a deep collection of quantitative data on nuclear division dynamics during the first three rounds of cell division in *C. elegans* embryos, in which 263 essential embryonic genes were silenced individually by RNA-mediated interference. The collection consists of datasets from 33 wild-type and 1142 RNAi-treated embryos, including five or more datasets for 189 genes. Application of a two-sample *t*-test identified 8660 reproducible RNAi-induced phenotypes for 421 phenotypic characters. Clustering analysis suggested 24 functional processes essential for early embryogenesis. Our collection is a rich resource for understanding animal development mechanisms.

**In Brief:** Kyoda et al. used bioimage informatics techniques to create a deep collection of quantitative data on nuclear division dynamics in RNAi-treated *C. elegans* embryos for 263 essential embryonic genes. Statistical analysis identified 8660 reproducible RNAi phenotypes for 421 phenotypic characters. The collection is a rich resource for understanding animal development.

**Highlights:**

- Bioimage informatics quantified nuclear division dynamics in *C. elegans* embryos
- From RNAi-silenced embryos we collected 1142 data sets on 263 essential genes
- Statistical analysis identified 8660 reproducible RNAi phenotypes
- Clustering analysis suggested 24 functional processes in *C. elegans* embryogenesis

## INTRODUCTION

A major challenge of developmental biology is to understand how molecular mechanisms generate multicellular structures such as bodies and organs from the fertilized egg. Recent progress in bioimage informatics techniques has enabled us to obtain quantitative data on multicellular dynamics from microscopy images of animal development. By using live-cell imaging and image-processing techniques, such quantitative data have been obtained for a wide variety of model organisms, ranging from *Caenorhabditis elegans* to mice (Bao et al., 2006; Bashar et al., 2012; Hamahashi et al., 2005; Keller et al., 2008, 2010).

One successful approach used to explore developmental molecular mechanisms has been the analysis of spatiotemporal cell-division dynamics when gene expression is perturbed (Neumann et al., 2010; Sönnichsen et al., 2005). In particular, several research groups have independently used bioimage informatics techniques in combination with differential interference contrast (DIC) microscopy or confocal microscopy imaging to create collections of quantitative data on nuclear division dynamics during *C. elegans* embryogenesis under a wide variety of gene silencing conditions (Ho et al., 2015; Kyoda et al., 2013; Santella et al., 2016). Such collections are practical for *C. elegans* because it is the only animal for which all essential embryonic genes have been identified through genome-wide RNA interference (RNAi) screens (Kamath et al., 2003; Sönnichsen et al., 2005). However, because only one or two sets of data are available for each condition, it is difficult to use commonly used statistical analysis methods such as a two-sample *t*-test for these collections.

A wild-type *C. elegans* embryo begins to generate a three-dimensional (3D) multicellular structure during the third round of cell division after fertilization (Figure 1A; Deppe et al., 1978; Sulston et al., 1983). The fertilized egg, P_0_, anteroposteriorly divides to produce a somatic founder cell, AB, and a germline cell, P_1_. AB divides dorsoventrally to produce two daughters, ABa and ABp, each of which divides in the left–right direction to produce two daughters: ABal and ABar, and ABpl and ABpr, respectively. P_1_ divides anteroposteriorly to produce a somatic cell, EMS, and a new germline cell, P_2_. EMS divides anteroposteriorly to produce two somatic founder cells, MS and E, whereas P_2_ divides dorsoventrally to produce a somatic founder cell, C, and a new germline cell, P_3_. Quantitative data on nuclear division dynamics in *C. elegans* embryos during the first three rounds of cell division will act as a basic resource for analyzing the mechanism generating 3D multicellular structures during animal development.

**Figure 1.**
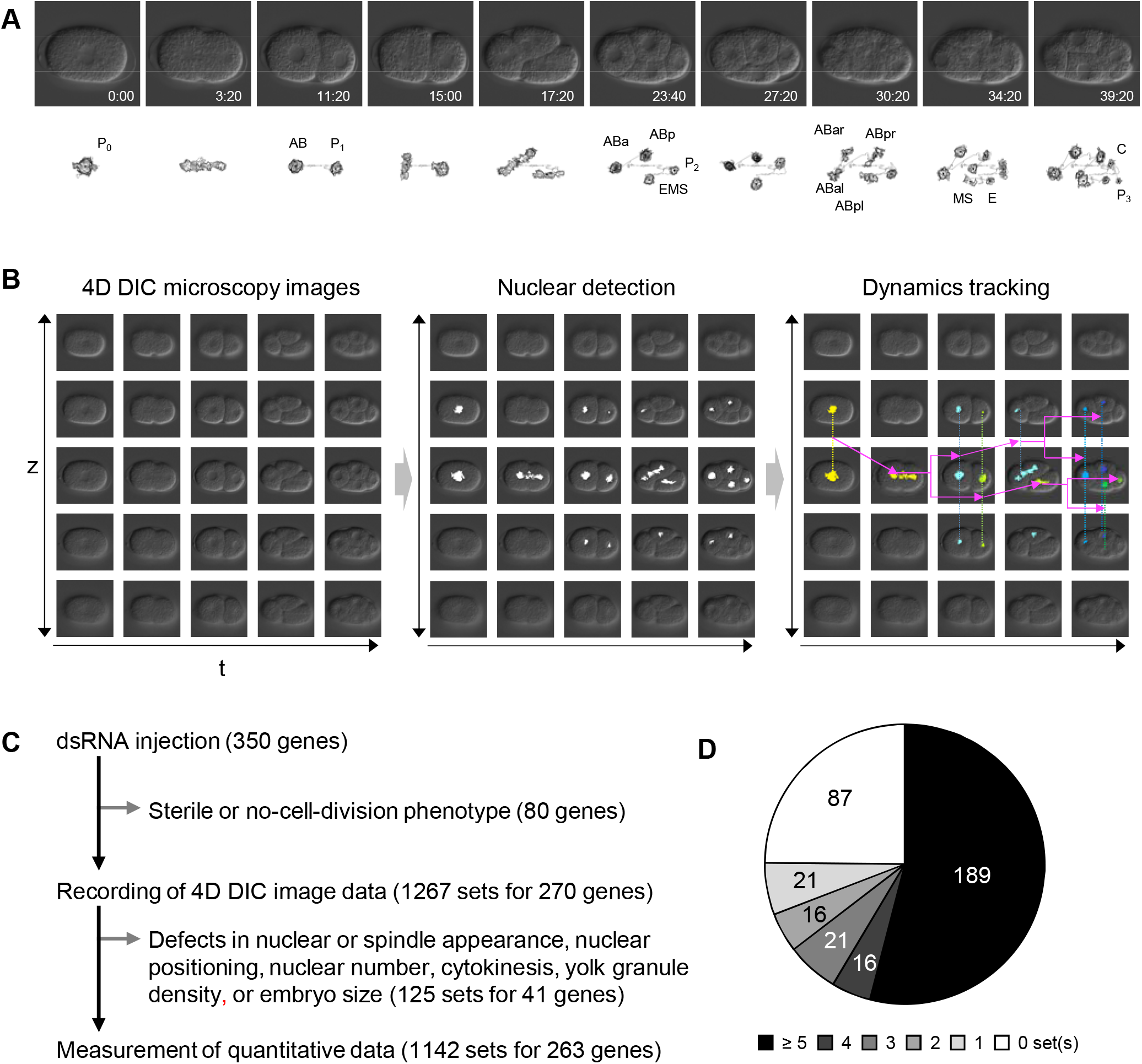
Collection of quantitative data on nuclear division dynamics in early *Caenorhabditis elegans* embryos. (A) Differential interference contrast (DIC) microscopy images (top) and 4D views of nuclear division dynamics data (bottom) in a wild-type embryo. P_0_, fertilized egg; AB and P_1_, anterior and posterior daughter cells, respectively, of P_0_; ABa and ABp, anterior and posterior daughter cells, respectively, of AB; ABxl and ABxr, left and right daughter cells, respectively, of ABx; EMS and P_2_, anterior and posterior daughter cells, respectively, of P_1_; MS and E, anterior and posterior daughter cells, respectively, of EMS; C and P_3_, dorsal and ventral daughter cells, respectively, of P_2_. (B) Overview of measurement of quantitative data on nuclear division dynamics from 4D (3D time-lapse) DIC microscopy image data. Different colored regions correspond to the nuclei in different cells. (C) Overview of the method used to obtain quantitative data on nuclear division dynamics from RNAi-treated embryos. (D) Distribution of the number of quantitative datasets for each gene.

Here, we created a deep collection of quantitative data on nuclear division dynamics in *C. elegans* embryos in which all 350 essential embryonic genes were silenced one at a time by using RNA-mediated interference (RNAi) (Fire et al., 1998). We obtained 1142 sets of quantitative data on 263 genes for the first three rounds of cell division. Because the collection included five sets of data for most of the genes examined, we could perform standard statistical analyses. We demonstrate that our collection provides a novel opportunity to understand the developmental molecular mechanisms of *C. elegans* embryos by performing computational analyses of the spatiotemporal nuclear division dynamics.

## RESULTS

### Deep Collection of Quantitative Data on Nuclear Division Dynamics

We first collected quantitative data on nuclear division dynamics from wild-type *C. elegans* embryos in the first three rounds of cell division. We recorded 3D time-lapse (4D) microscopy images of wild-type embryos by using Nomarski DIC microscopy (Figure 1A). We then detected the outlines of nuclei and spindles in a 3D coordinate system and tracked them by using our automated system (Figure 1B; Hamahashi et al., 2007). We obtained 33 sets of quantitative data on nuclear division dynamics in wildtype embryos (Figure 1A).

We next obtained a deep collection of quantitative data on nuclear division dynamics in gene-perturbed embryos. We used RNAi to silence, individually, all 350 genes for which an embryonic lethal phenotype had been produced in 100% of offspring in a previous genome-wide RNAi screen (Figure 1C; Kamath et al., 2003). For each of the 350 essential genes, we injected the corresponding double-stranded (ds)RNA into between 5 to 24 young adult worms with the aim of obtaining five or more sets of quantitative data from different RNAi-treated embryos. For 270 genes, we obtained 1267 sets of 4D DIC microscopy images in which RNAi-treated embryos underwent cell division (Figure 1C). For 225 of the 270 genes, we obtained five or more sets of images per gene. However, for the other 45 genes, we obtained between one and four sets per gene, because more than half of the RNAi-treated worms produced no embryos (sterile phenotype) or more than half of the RNAi-treated embryos did not undergo cell division (no-cell-division phenotype). No images were obtained for the other 80 of the 350 genes, because all the RNAi-treated worms or embryos exhibited the sterile or nocell-division phenotype (Figure 1C). We then applied our automated system to the 1267 image datasets. For 125 image datasets corresponding to 41 genes, we could not obtain quantitative data because the RNAi-treated embryos exhibited abnormalities in nuclear appearance, spindle appearance, nuclear positioning (two nuclei were in close proximity at the cortex), nuclear number, cytokinesis (including multiple nuclei), yolk granule density, or embryo size (large). Our automated system depends on the nuclear and spindle appearance, the separable nuclear positioning, the difference between nuclear and cytoplasmic image texture, the embryo size, and the assumption that only one nucleus exists in a cell and the nuclei do not fuse (Hamahashi et al., 2007). From RNAi-treated embryos we obtained 1142 sets of quantitative data for 263 genes (Figure 1C)— 75% of the 350 essential embryonic genes (Figure 1D). We obtained five or more sets of quantitative data for 189 genes (54% of the 350 genes) (Figure 1D). The resultant datasets are available online at Worm Developmental Dynamics Database 2 (WDDD2; https://wddd.riken.jp).

In our experiments, RNAi of 29% (102/350) of the 350 genes resulted in 100% embryonic lethality, 60% (209/350) in partial embryonic lethality, and 11% (39/350) in no embryonic lethality (Table S1).

### Analysis of Cell-division Dynamics in Wild-type Embryos

To evaluate the quality of the quantitative data obtained, we analyzed the nuclear division patterns of the wild-type embryos in our collection. The nuclear division patterns of wild-type *C. elegans* embryos are largely invariant (Sulston et al., 1983). To verify the patterns in our collection, we first analyzed the order of cell division. The nucleus is spherical during the interphase; it becomes less spherical when it enters the mitotic phase because of nuclear envelope breakdown and the growth of centrosomal microtubules at the nuclear poles. After cell division, the daughter nuclei become spherical again when they reform nuclear envelopes and enter interphase (Figure 2A). Therefore, we determined the division timing for each cell by calculating the sphericity of the nuclear regions (see Star Methods). In wild-type *C. elegans* embryos, the cells divide in the order of P_0_, AB, P_1_, ABa and ABp (synchronous), EMS, and P_2_ (Figure 1A; Deppe et al., 1978; Sulston et al., 1983) during the first three rounds of cell division. We confirmed that the cell divisions in the wild-type embryos in our collection were consistent with the canonical order (Figure 2B).

**Figure 2.**
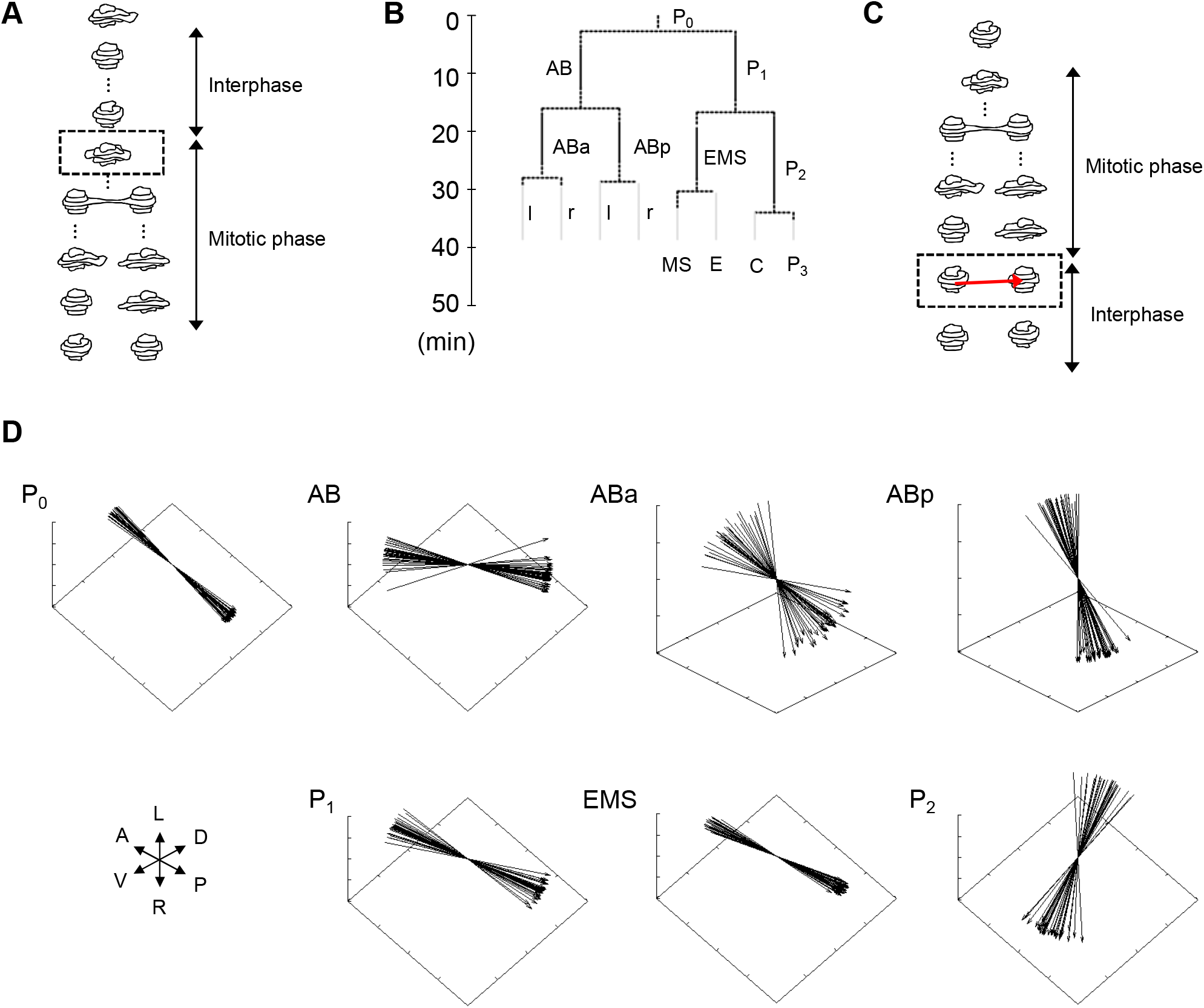
Computational analysis of nuclear division dynamics in wild-type embryos. (A) Method used to determine the duration of cell division. Each object represents a nuclear region. Time progresses from top to bottom. The dashed square represents the moment in cell division when the sphericity of the nuclear region becomes less than the predefined threshold for the first time. (B) Nuclear division pattern obtained from wildtype embryo data in the collection. Cell names are defined as in Figure 1. l and r represent ABxl and ABxr, respectively. (C) Method used to determine the orientation of cell division. The red arrow represents the orientation of cell division, as calculated from the positions of the two daughter cells’ nuclear regions, and the dashed square represents the moment in the calculation when the sphericities of the nuclear regions of both daughter cells exceed the predefined threshold for the first time. (D) Orientation of cell divisions from the wild-type embryo data in the collection. A, P, D, V, L, and R represent anterior, posterior, dorsal, ventral, left, and right, respectively.

We next analyzed the orientations of the cell divisions. We determined the orientation of a cell division by calculating a vector between two daughter cells’ nuclei just after mitotic phase (Figure 2C). In wild-type embryos, P_0_, P_1_, and EMS divide along the anterior–posterior (AP) axis, AB and P_2_ divide along the dorsal–ventral (DV) axis, and ABa and ABp divide along the left–right (LR) axis (Deppe et al., 1978; Sulston et al., 1983). We found that the orientations of the division axes of all cells in our collection were consistent with the canonical orientations (Figure 2D). These results indicated that our collection was of sufficient quality for phenotypic analysis of celldivision dynamics.

### RNAi-induced Phenotypes Detected by Two-sample *t*-test

To demonstrate that our collection provided a rich resource for phenotypic analysis, we computationally detected RNAi-induced phenotypes from the collection. We focused on the RNAi-induced phenotypes related to nuclear division dynamics and reported as the 16 defect categories in a previous manual analysis of pronuclear fusion to the four-cell stage (Sönnichsen et al., 2005). We mathematically defined 51 phenotypic characters that could mimic these 16 defect categories in cells from pronuclear fusion to the four-cell stage (Table S2). We then extended these characters to cells from the four-cell stage to the eight-cell stage. In total, we defined 421 phenotypic characters that indexed the position, size, and shape of the nucleus and spindle; the arrangement of cells; the orientation of cell-division axes; and the timing of cell divisions (Table S3). By extracting the values of these characters from each dataset, we obtained phenotypic expression profiles of wild-type and RNAi-treated embryos (Table S4).

To detect RNAi-induced phenotypes, we statistically compared the values for each character between wild-type and RNAi-treated embryos. We first applied a method based on a two-sample unequal variance *t*-test (Welch’s *t*-test) commonly used in experimental biology (Ruxton, 2006), followed by multiple test correction (Storey and Tibshirani, 2003). Statistically significant differences between wild-type and RNAi-treated embryos were classed as RNAi-induced phenotypes. Among 263 genes for 421 phenotypic characters, we detected 8660 RNAi-induced phenotypes from the phenotypic expression profiles (Table S5).

To validate the results of our phenotypic analysis, we examined whether our analysis had detected the RNAi-induced phenotypes reported in a previous large-scale manual analysis (Sönnichsen et al., 2005). For the 263 genes tested in our analysis, the previous analysis reported 203 RNAi-induced phenotypes for the 16 defect categories (Table 1). Out of these 203 phenotypes, 149 were observable in at least one set of our DIC microscopy images. Our method detected about 68% (102/149) of the previously reported phenotypes that were also reproduced in our RNAi experiments. The remaining 47 phenotypes reproduced in our experiments were not detected. For 21 of these phenotypes, the number of RNAi datasets was too small (<2) to apply the two-sample *t*-test. For the other 26 phenotypes, the penetrance or expressivity, or both, of the RNAi silencing were low (e.g., one of the five RNAi datasets exhibited the phenotype), so the test’s statistical power was insufficient to detect the phenotype.

**Table 1.** Comparison of RNAi-induced phenotypes detected in our study with those reported in a previous manual analysis (Sonnichsen et al., 2005)

| # | Defect category | Share | Reproduced | Two-sample *t*-test | Data transformation | Combined | New |
| --- | --- | --- | --- | --- | --- | --- | --- |
| 1 | P<sub>0</sub> pronuclear centration | 10 | 6 | 3 | 4 | 5 | 51 |
| 2 | P<sub>0</sub> pronuclear rotation | 4 | 3 | 0 | 3 | 3 | 145 |
| 3 | P<sub>0</sub> spindle integrity | 13 | 10 | 6 | 10 | 10 | 243 |
| 4 | P<sub>0</sub> spindle elongation | 4 | 4 | 0 | 4 | 4 | 156 |
| 5 | P<sub>0</sub> spindle positioning | 13 | 7 | 5 | 7 | 7 | 82 |
| 6 | P<sub>1</sub>/AB nuclear separation | 19 | 9 | 4 | 7 | 9 | 119 |
| 7 | AB nuclear migration | 2 | 0 | 0 | 0 | 0 | 64 |
| 8 | P<sub>1</sub>/AB nuclei size/shape | 25 | 19 | 13 | 17 | 19 | 200 |
| 9 | P<sub>1</sub> nuclear migration/rotation | 6 | 5 | 2 | 5 | 5 | 121 |
| 10 | P<sub>1</sub>/AB spindle assembly | 4 | 4 | 2 | 3 | 3 | 233 |
| 11 | AB spindle orientation | 3 | 4 | 2 | 3 | 3 | 122 |
| 12 | P<sub>1</sub>/AB asynchrony of divisions | 32 | 27 | 20 | 26 | 27 | 147 |
| 13 | Four-cell-stage cross-eyed | 18 | 13 | 11 | 12 | 13 | 160 |
| 14 | Four-cell-stage nuclei size/shape | 34 | 27 | 24 | 27 | 27 | 235 |
| 15 | Four-cell-stage configuration | 4 | 4 | 3 | 4 | 4 | 59 |
| 16 | Overall pace of events | 12 | 9 | 7 | 9 | 9 | 193 |
| Total |  | 203 | 149 | 102 | 142 | 148 | 2330 |

Share: number of genes shared by a previous manual analysis (Sönnichsen et al., 2005) and our analysis. Reproduced: number of phenotypes reproduced in our RNAi experiments. Two-sample *t*-test: number of phenotypes reproduced in our RNAi experiments and detected by using a method based on the two-sample *t*-test. Data transformation: number of phenotypes reproduced in our RNAi experiments and detected by using a method based on data transformation and *P*-value estimation. Combined: number of phenotypes reproduced in our RNAi experiments and detected by using one or both of the two methods. New: number of phenotypes that were not detected by the previous analysis but were detected in our study by using one or both of the two methods.

### RNAi-induced Phenotypes Detected by Data Transformation and *P*-value Estimation

To overcome the problems of sample size and penetrance and expressivity of RNAi silencing, we developed a new method based on data transformation and *P*-value estimation (see Star Methods). In this method, we first transformed the values for each phenotypic character extracted from the wild-type embryos to follow a normal distribution. Next, we transformed the values of each phenotypic character extracted from RNAi-treated embryos by using the same data transformation parameters as used for the wild-type embryos. We then estimated the *P*-values of the values from RNAi-treated embryos on the basis of the corresponding normal distributions. The method can be used to detect RNAi-induced phenotypes when at least one phenotypic alteration appears in the RNAi-treated embryos.

By applying the method to the phenotypic expression profiles, we detected 21,034 RNAi-induced phenotypes for the 263 genes. These phenotypes were derived from 70,845 alterations detected in 1142 RNAi-treated embryos (Table S6). The method based on data transformation and *P*-value estimation detected about 95% (142/149) of the previously reported phenotypes reproduced in our RNAi experiments (Table 1). Thus, the detection sensitivity of this method (95%) is higher than that of the method based on a two-sample *t*-test (68%). We were able to detect phenotypes for genes for which only one dataset was available or that had low repeatability. These results suggest that the method based on data transformation and *P*-value estimation can be of practical use in the case of limited datasets and low penetrance or expressivity of phenotypic alterations.

### RNAi-induced Phenotypes Detected by the Two Statistical Methods

We detected a total of 26,539 RNAi-induced phenotypes for 421 phenotypic characters from the datasets for 263 genes by using the two statistical methods. The method based on the two-sample *t*-test and that based on data transformation and *P*-value estimation detected 8660 and 21,034 RNAi-induced phenotypes, respectively (Figure 3A), with 3155 phenotypes being detected by both methods. Because the two methods detected different types of RNAi-induced phenotypes, the methods appeared to be complementary.

**Figure 3.**
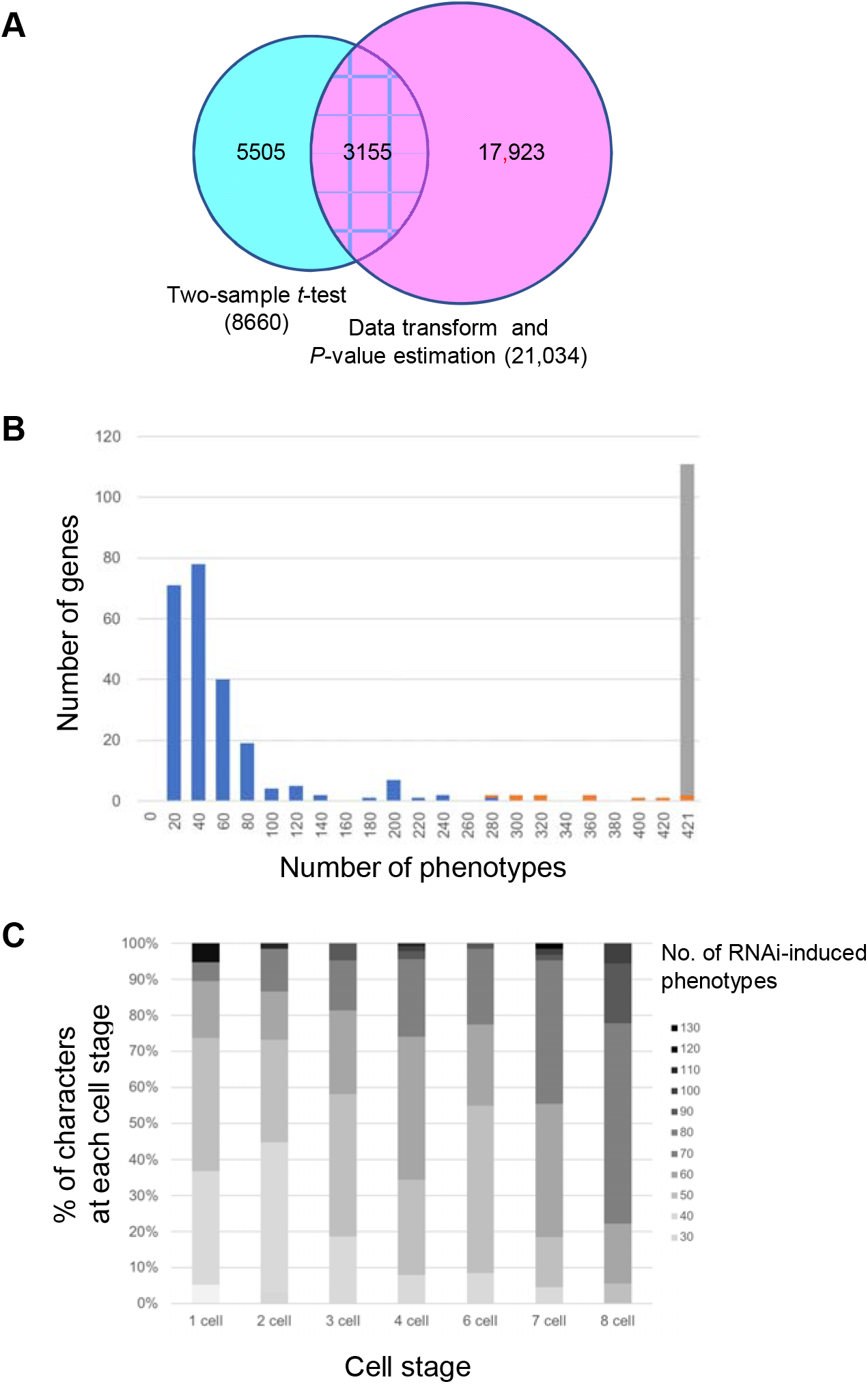
Computational analysis of RNAi-induced phenotypes. (A) Venn diagram of the numbers of RNAi-induced phenotypes detected by the method based on the two-sample *t*-test and that based on data transformation and *P*-value estimation. (B) Histogram of the number of genes with the indicated numbers of RNAi-induced phenotypes among the 421 mathematically defined phenotypic characters. Bins 0, 20, …, 420, 421 represent 0, 1 to 20, … , 401 to 420, 421, respectively. Blue, orange, and gray represent the genes for which we could calculate more than half, fewer than half, and none, respectively, of the 421 phenotypic characters in the RNAi-treated embryos. (C) Distribution of the percentage and number of RNAi-induced phenotypes for each phenotypic character at each cell stage (one-cell to eight-cell). RNAi-induced phenotypes detected by a method based on the two-sample *t*-test were used (B, C).

We combined the results of the two statistical methods and compared them with those of a previous large-scale manual analysis (Sönnichsen et al., 2005). We found that the two methods, when combined, detected all the phenotypes except one for the 16 defect categories reported in the previous manual analysis and reproduced in our RNAi experiments (Table 1). This result indicates the validity of our computational phenotypic analysis using phenotypic expression profiles.

We next focused on new phenotypes detected for the 16 defect categories. Our computational analyses detected 2478 phenotypes for the 16 defect categories in total. Among them, 148 phenotypes were reported in the previous manual analysis (Sönnichsen et al., 2005). Thus, 94% (2330/2478) of the phenotypes were new phenotypes found in our analyses (Table 1). Statistical analysis could have detected subtle but significant phenotypic alterations that were difficult to detect by manual analysis. Some RNAi-induced phenotypic alterations may have occurred only in our experiment. On the basis of the 94% value, we estimated that 24,954 RNAi-induced phenotypes were new phenotypes found for the 421 phenotypic characters in our analysis.

### Features of RNAi-induced Phenotypes in Early *C. elegans* Embryogenesis

To explore the variations in phenotypic expression among the 263 genes, we calculated the number of RNAi-induced phenotypes for each gene and then compiled the distribution of these numbers (Figure 3B; Figure S1A). Phenotypic characters that differed significantly between wild-type and RNAi-treated embryos were classed as RNAi-induced phenotypes. Phenotypic characters that could not be calculated because of severe defects were also classed as RNAi-induced phenotypes. We found that all 263 essential embryonic genes exhibited at least one phenotype in nuclear division dynamics (Figure 3B). This finding indicates that all the essential embryonic genes are involved in developmental processes during the first three rounds of cell division, or earlier.

We also found that the histogram had two peaks: a) a broad peak with a right skew on the left side, corresponding to low numbers of phenotypes (1 to 80 phenotypes), and b) a narrow peak on the right side, corresponding to all 421 phenotypes examined (Figure 3B). All genes counted in the left-hand peak led to some defects in cell division but almost normal development, because we did not detect RNAi-induced phenotypes for most phenotypic characters in these genes (Figure 3B). The shape of the histogram on the left side was similar to that obtained from the results of a large-scale phenotypic analysis of non-essential genes in *Saccharomyces cerevisiae* (Ohya et al., 2005). This result suggests that the genes counted in this peak are dispensable for mitosis but still essential for embryogenesis. RNAi of the genes counted in the narrow peak on the right side of the histogram resulted in extremely severe defects: in almost all cases, no phenotypic characters could be calculated in these embryos (Figure 3B; Table S7). By using gene ontology (GO) enrichment analysis (Mi et al., 2013), we found that the genes counted in this peak were essential for cell-cycle processes, including both meiosis and mitosis (Table S7). In the vicinity of the right narrow peak we also found enrichment of genes specific to the mitotic cell cycle (Figure 3B; Table S7). RNAi of these genes would cause severe defects in one-cell-stage embryos. Fewer than half of the phenotypic characters could be calculated in these embryos.

To investigate whether or not the effect of RNAi was cumulative, we calculated the number of RNAi-induced phenotypes for each phenotypic character and then compiled the distribution of these numbers at each cell stage (Figure 3C; Figure S1B). We found that the percentage of phenotypic characters exhibiting larger numbers of RNAi-induced phenotypes increased over time. This result suggests that the effect of RNAi is cumulative: phenotypic defects at an earlier cell stage affect the events at later cell stages.

### Phenotypes Pertaining to Spatiotemporal Dynamics of Cell Division

Our collection enables us to detect temporal and 3D spatial phenotypes at a single-cell level. An example of a temporal phenotypic character is the duration of cell-cycle phases. We determined the duration of cell-cycle phases for each cell by calculating the sphericity of the nuclear regions (Figure 2A; see Star Methods). We identified 497 RNAi-induced phenotypes for the duration of cell-cycle phases for the 263 genes; examples are shown in Figure 4A. The first three examples demonstrate different types of cell-cycle-phase-specific RNAi-induced phenotypes. The gene *pola-1* encodes an ortholog of human POLA1 (DNA polymerase alpha 1, catalytic subunit) (Kim et al., 2018) and is predicted to have DNA binding activity, DNA-directed DNA polymerase activity, and nucleotide-binding activity (Camon et al., 2005). The previous RNAi screen reported delayed pronuclear rotation, off-centric AB migration, and late P_1_ division in its RNAi embryos (Sönnichsen et al., 2005). We found that *pola-1*(*RNAi*) embryos exhibited longer interphases and mitotic phases than in the wild type in many cells during the first three rounds of cell division (67% (4/6) of interphases and 86% (6/7) of mitotic phases). *pola-1*(*RNAi*) embryos demonstrate an RNAi-induced phenotype that affects both interphases and mitotic phases. The gene *ucr-1* encodes an ortholog of human UQCRC1 (ubiquinol-cytochrome reductase core protein 1) (Kim et al., 2018). The previous RNAi screen reported a slow pace of progression from pronuclear meeting to AB cell division (Sönnichsen et al., 2005). We found that *ucr-1*(*RNAi*) embryos exhibited longer interphases in all cells during the first three rounds of cell division. No significant difference compared with the wild type was found in the duration of mitotic phases. Thus *ucr-1(RNAi)* embryos demonstrate an interphase-specific RNAi-induced phenotype. The gene *lin-53* encodes an ortholog of human RBBP4 (RB binding protein 4, chromatin remodeling factor) and RBBP7 (RB binding protein 7, chromatin remodeling factor) (Kim et al., 2018) and exhibits histone deacetylase binding activity (Lu and Horvitz, 1998). The previous RNAi screen reported no significantly different phenotype in early embryogenesis compared with the wild type (Sönnichsen et al., 2005). Unlike *ucr-1(RNAi)* embryos, we found that, compared with the wild type, *lin-53*(*RNAi*) embryos exhibited longer mitotic phases in all cell divisions except P_2_ division during the first three rounds of cell division (Figure 4A). No significant difference was found in the duration of interphase except ABp interphase. Thus *lin-53*(*RNAi*) embryos demonstrate a mitotic-phase-specific RNAi-induced phenotype. The latter two examples respectively demonstrate cell-stage-specific and lineage-specific RNAi-induced phenotypes. The gene *teg-4* encodes an ortholog of human SF3B3 (splicing factor 3b subunit 3) (Kim et al., 2018) and is predicted to have nucleic acid binding activity (Camon et al., 2005). The previous RNAi screen reported no significantly different phenotype compared with the wild type in early embryogenesis (Sönnichsen et al., 2005). We found that *teg-4(RNAi)* embryos exhibited longer interphases, but only in all cells at the four-cell stage. Thus *teg-4(RNAi)* embryos demonstrate a cell-stage-specific RNAi-induced phenotype. The gene *gei-4* is involved in negative regulation of Ras protein signal transduction and vulval development (Poulin et al., 2005). The previous RNAi screen reported no significantly different phenotype compared with the wild type in early embryogenesis (Sönnichsen et al., 2005). We found that *gei-4(RNAi)* embryos exhibited longer interphases, but only in all cells in the P lineage; that is, P_1_ and P_2_. Thus *gei-4*(*RNAi*) embryos demonstrate a cell-lineage-specific RNAi-induced phenotype. Our collection therefore enables us to detect RNAi-induced temporal phenotypes specific to cell-cycle phases, cell stages, and cell lineages.

**Figure 4.**
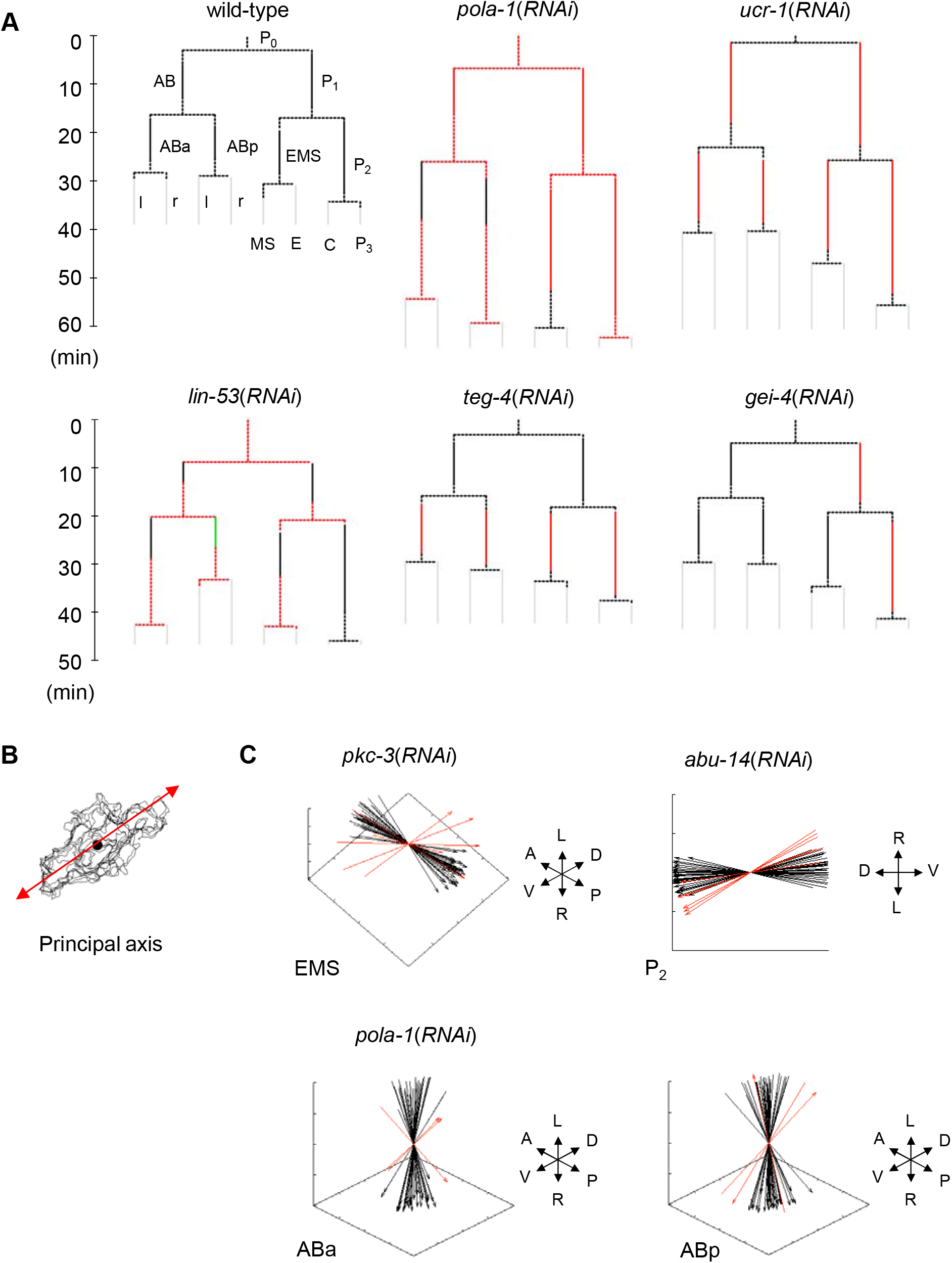
Spatiotemporal phenotypes detected in our computational phenotypic analysis. (A) Examples of abnormal division timing from *pola-1*(*RNAi*), *ucr-1*(*RNAi*), *lin-53*(*RNAi*), *teg-4*(*RNAi*), and *gei-4*(*RNAi*) embryos. For comparison, the average cellcycle length in wild-type embryos is shown (top left). Cell names are defined as in Figure 1. Solid lines represent interphase and dotted lines represent mitotic phase. Cellcycle phases elongated, shortened, or unchanged in RNAi-treated embryos are shown in red, green, or black, respectively. Gray lines represent the interphases that were not measured. (B) Method used to determine spindle orientation. (C) Examples of abnormal spindle orientation from*pck-3*(*RNAi*), *abu-14*(*RNAi*), and *pola-1*(*RNAi*) embryos. Spindle orientations of RNAi-treated embryos are shown by red arrows and those of wild-type embryos are shown by black arrows.

An example of a 3D spatial phenotypic character is the spindle orientation. We determined the orientation by calculating the first principal axis of the nuclear region at the beginning of the mitotic phase (Figure 4B). We detected 111 RNAi-induced phenotypes of spindle orientation for the 263 genes; examples are shown in Figure 4C. The following three examples demonstrate RNAi-induced spindle orientation phenotypes in different orientation axes. The gene *pkc-3* encodes an ortholog of human PRKCI (protein kinase C iota) and PRKCZ (protein kinase C zeta) (Kim et al., 2018). In wild-type embryos, PKC-3 protein co-localizes with PAR-3 protein, which regulates cell polarity, and localizes between ABa and EMS cells (Tabuse et al., 1998). We found that the spindle orientations in EMS were tilted slightly toward the DV axis in *pkc-3(RNAi)* embryos, whereas the orientation was along the AP axis in wild-type embryos. Thus *pkc-3*(*RNAi*) embryos demonstrate an RNAi-induced spindle orientation phenotype in AP-oriented spindles. The gene *abu-14* is involved in pharynx development (George-Raizen et al., 2014). A previous RNAi screen reported a 100% embryonic lethal phenotype (Kamath et al., 2003), but no detailed phenotypes in embryos have been reported. We found that the spindles in P_2_ were tilted slightly toward the LR axis in the *abu-14(RNAi)* embryo, whereas the orientation was along the DV axis in wild-type embryos. Thus *abu-14(RNAi)* embryos demonstrate an RNAi-induced spindle orientation phenotype in DV-oriented spindles. The gene *pola-1* is an ortholog of human POLA1 (DNA polymerase alpha 1, catalytic subunit). The previous RNAi screen (Kamath et al., 2003) reported delayed pronuclear rotation, off-centric AB migration, and late P_1_ division. We found that the spindle orientations were random in ABa and ABp in *pola-1*(*RNAi*) embryos, whereas the orientation was along the LR axis in wild-type embryos. Thus *pola-1*(*RNAi*) embryos demonstrate an RNAi-induced spindle orientation phenotype in LR-oriented spindles. Our collection therefore enables us to detect RNAi-induced 3D spatial phenotypes for any axial directions.

### Clustering of Phenotype Expression Profiles

To determine which biological processes are required for nuclear division dynamics during the first three rounds of cell division, we performed a clustering analysis using our phenotypic expression profiles (see Star Methods). In this analysis, we assumed that silencing of functionally associated genes would result in similar phenotypic expression patterns. By using hierarchical clustering, 1141 of the 1142 RNAi-treated embryos were classified into 24 clusters (Figure 5A; Table S8); a *lam-2(RNAi)* embryo was excluded as an outlier (see Star Methods). The clusters were determined manually from the tree structure and heat map of the clustering result.

**Figure 5.**
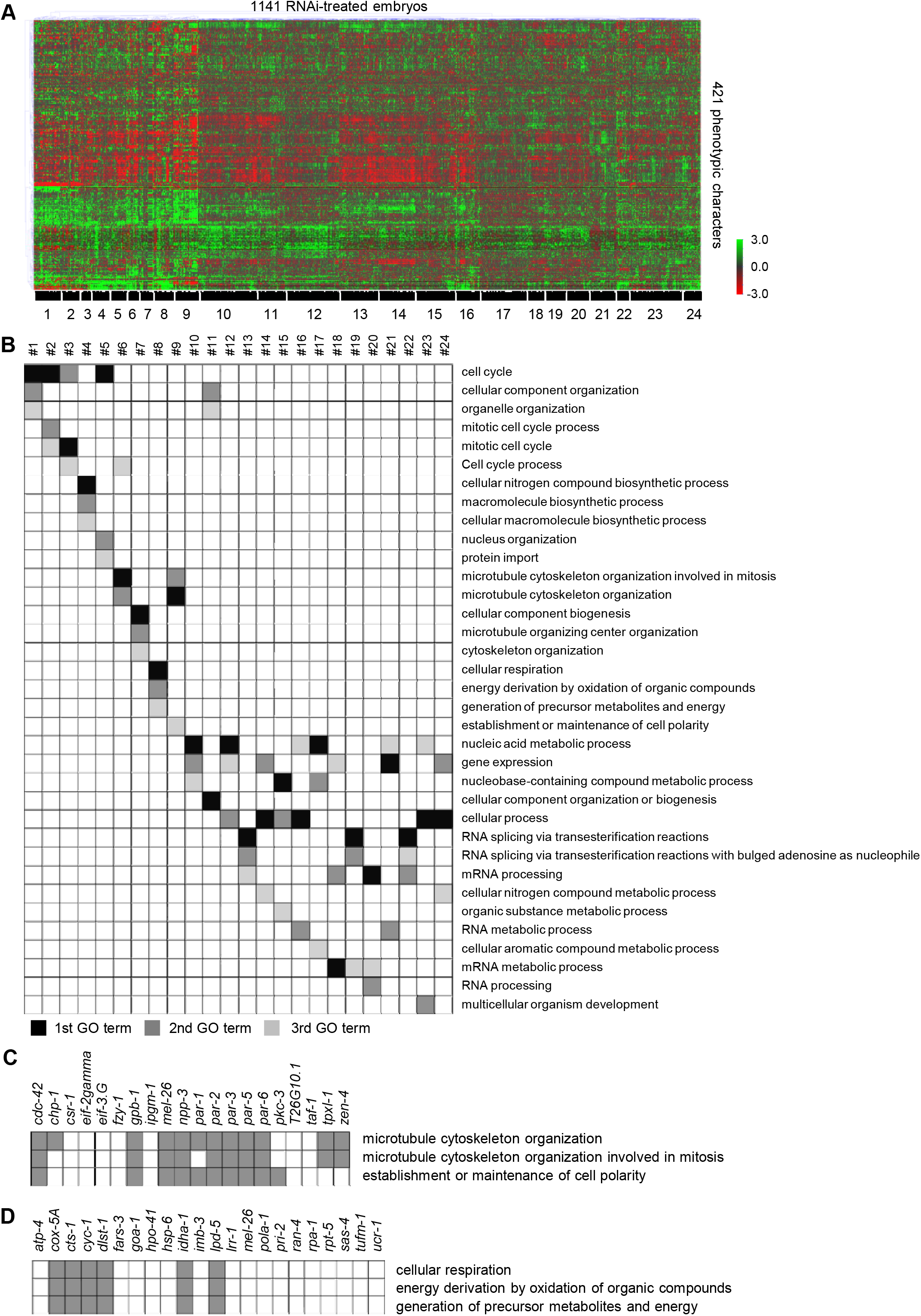
Clustering of phenotypic expression profiles. (A) Hierarchical clustering of the phenotypic expression profiles for 1141 RNAi-treated embryos and 421 phenotypic characters. The 1141 RNAi-treated embryos could be separated into 24 gene clusters (black bars). These clusters were determined manually by using information from the hierarchical tree structure and the heat map. (B) Gene ontology (GO) enrichment analysis for the 24 gene clusters yielded a heatmap of the top three Biological Process GO terms for each of the 24 clusters. Black, mid-gray, and light gray represent the first, second, and third significant GO terms, respectively. (C, D) Table of correspondence of annotations of the top three Biological Process GO terms to the genes in cluster 9 (C) and cluster 8 (D).

To generate functional annotation of the clusters, we performed GO enrichment analysis of the genes in each cluster. Because many of the top Biological Process GO terms with the smallest false-discovery-rate (FDR)-corrected *P*-values were shared among several clusters, we annotated each cluster by using the top three Biological Process GO terms (Figure 5B; Table S9). By using the top three GO terms, all clusters were annotated differently (Figure 5B).

To demonstrate how these three GO terms represented the functions of genes in each cluster, we explored two clusters as examples. Cluster 9 consisted of 20 genes (Figure 5C) and was annotated with “microtubule cytoskeleton organization,”“microtubule cytoskeleton organization involved in mitosis,” and “establishment or maintenance of cell polarity” (Figure 5B). Eight of the 20 genes were annotated with all the GO terms, whereas 13 were annotated with one or more of the three GO terms (Figure 5C; Table S10). Among the seven genes that were not annotated with any of the three GO terms, two, namely *fzy-1* and *csr-1,* had functions associated with the three GO terms (Table S10); *fzy-1* is involved in polarity specification of the anterior–posterior axis (Rappleye et al., 2002), and *csr-1* is involved in P-granule assembly (Andralojc et al., 2017). P-granules are RNA–protein condensates in the germline of *C. elegans* and are asymmetrically localized in the cytoplasm of newly fertilized embryos (Seydoux, 2018). The cluster also contained genes involved in translational initiation factor activity (*eif-2gamma* and *eif-3.G*), transcriptional regulation activity (*taf-1*), and glucose catabolic process (*ipgm-1*) as well as an uncharacterized gene (T26G10.1) (Table S10). Cluster 9 likely corresponds to a functional process that is well represented by the annotated three GO terms, but it likely includes cellular processes beyond these terms.

Cluster 8 consisted of 22 genes (Figure 5D) and was annotated with “cellular respiration,”“energy derivation by oxidation of organic compounds,” and “generation of precursor metabolites and energy” (Figure 5B). Six out of the 22 genes were annotated with all three GO terms (Figure 5D; Table S10). Among the 16 genes not annotated with the three GO terms, six—*atp-4, hsp-6, tufm-1, ucr-1, fars-3,* and *rpt-5—*have functions associated with the three GO terms. Specifically, four of them—*atp-4, hsp-6, tufm-1,* and *ucr-1—*are expressed in the mitochondria, which play the central role in cellular respiration (Mitchell, 1961). The remaining two genes, *fars-3* and *rpt-5,* are involved in tRNA and the amino acid metabolic process, respectively (Table S10). The cluster also contains genes involved in DNA-dependent DNA replication (*pola-1, pri-2,*and *rpa-1*), protein import into the nucleus (*imb-3* and *ran-4*), the ubiquitylation system (*lrr-1* and *mel-26*), the G-protein-coupled receptor signaling pathway (*goa-1*), and the centrosome cycle (*sas-4*) (Table S10). *hpo-41* is a pseudogene. Cluster 8 likely represents a group of functions that is spread widely across many cellular processes, including those represented by the three annotated GO terms.

### Annotation of Uncharacterized Genes

Among the 263 genes for which 1141 of the 1142 RNAi-treated embryos were analyzed, 18 were uncharacterized: that is, they were not given approved gene names consisting of a three- or four-letter gene-class name followed by a hyphen and a number in the WormBase database (https://wormbase.org; WS277 release). To predict the functions of these 18 genes, we collected the GO terms annotated to the clusters containing each of them. The 18 genes were annotated with 62 sets of three GO terms; each three-GO-term set corresponded to a single cluster (Table S11). Many genes belonged to more than one cluster. We predicted the functions of the 18 genes by using the GO terms collected for each of them (Table 2).

**Table 2.**
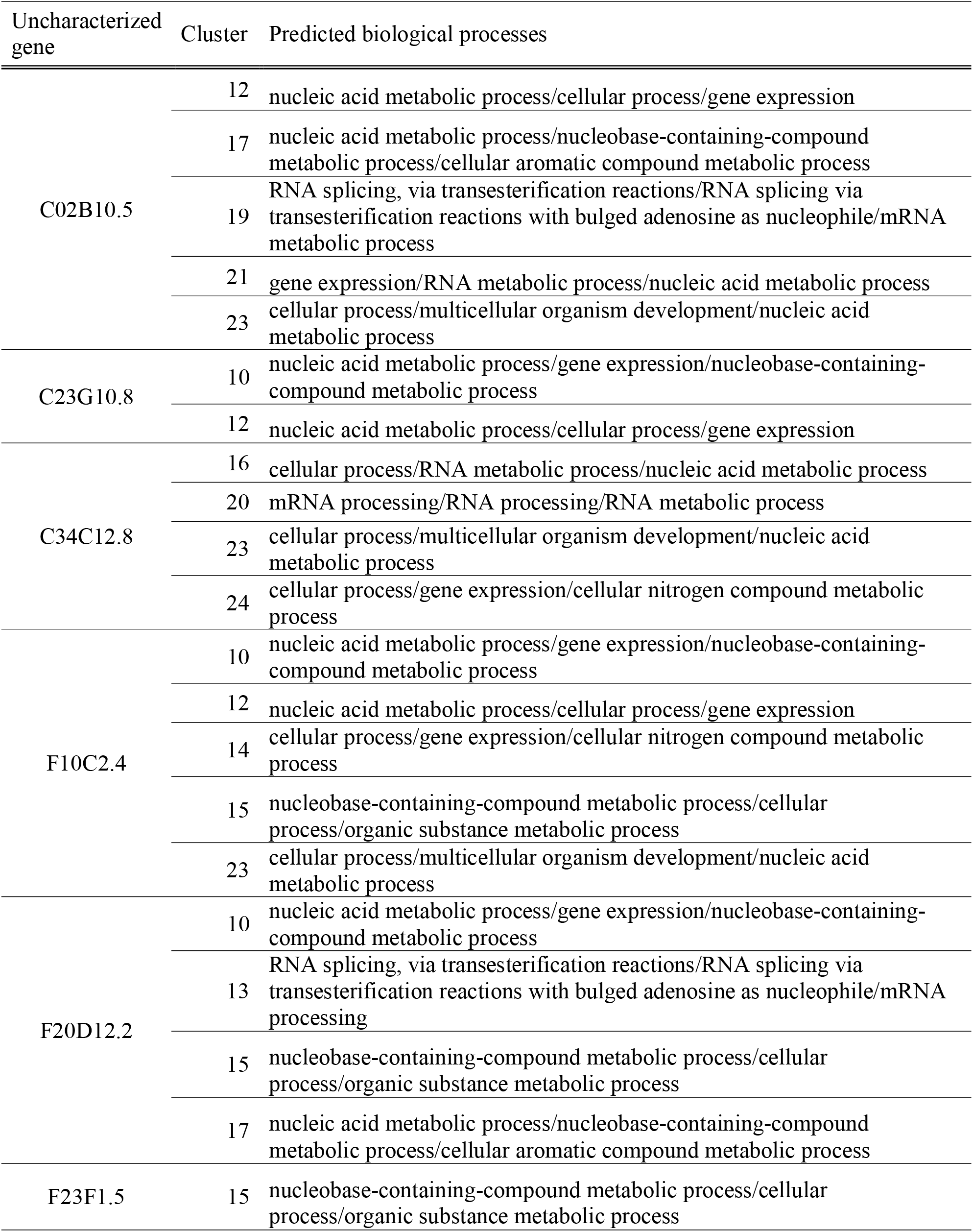
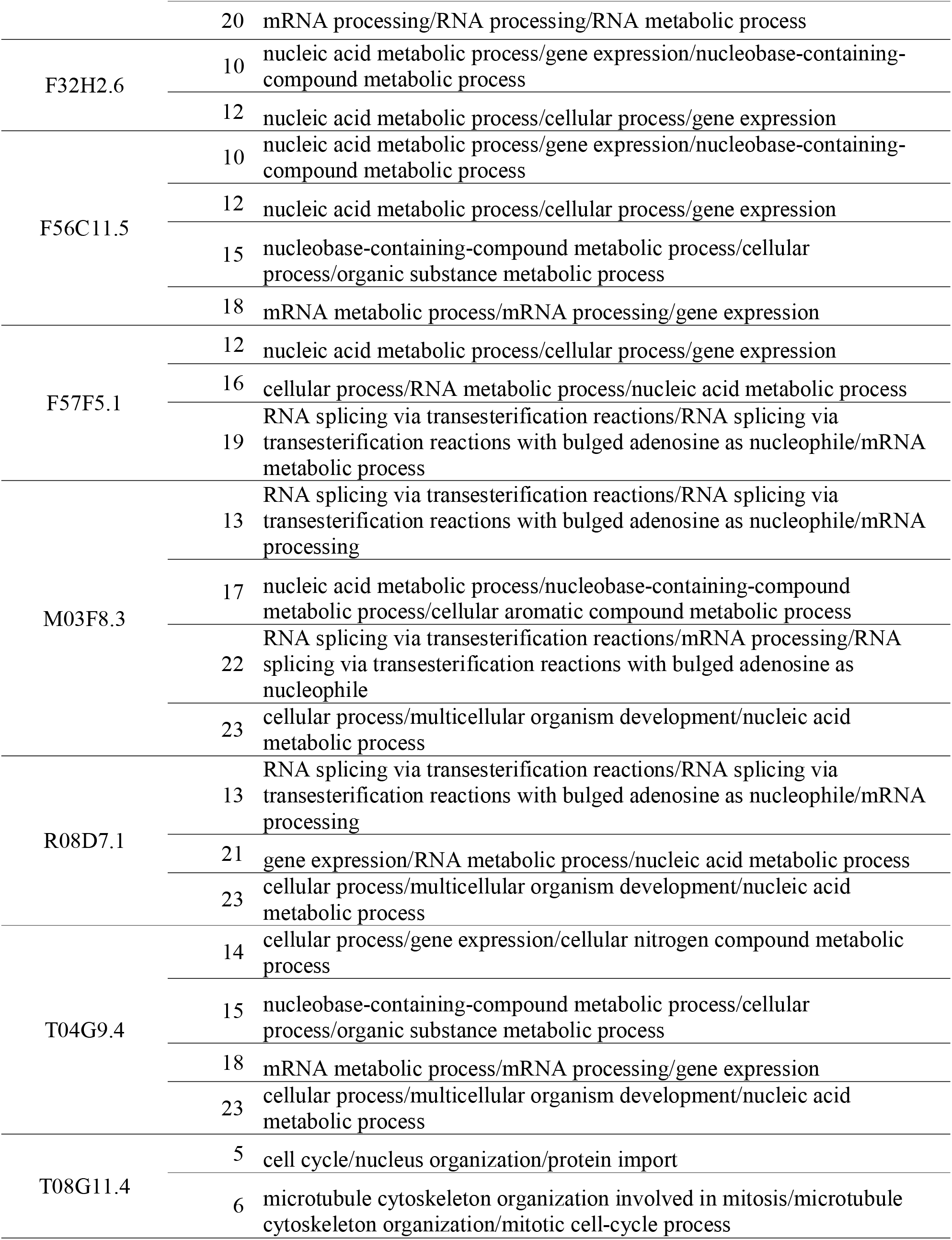
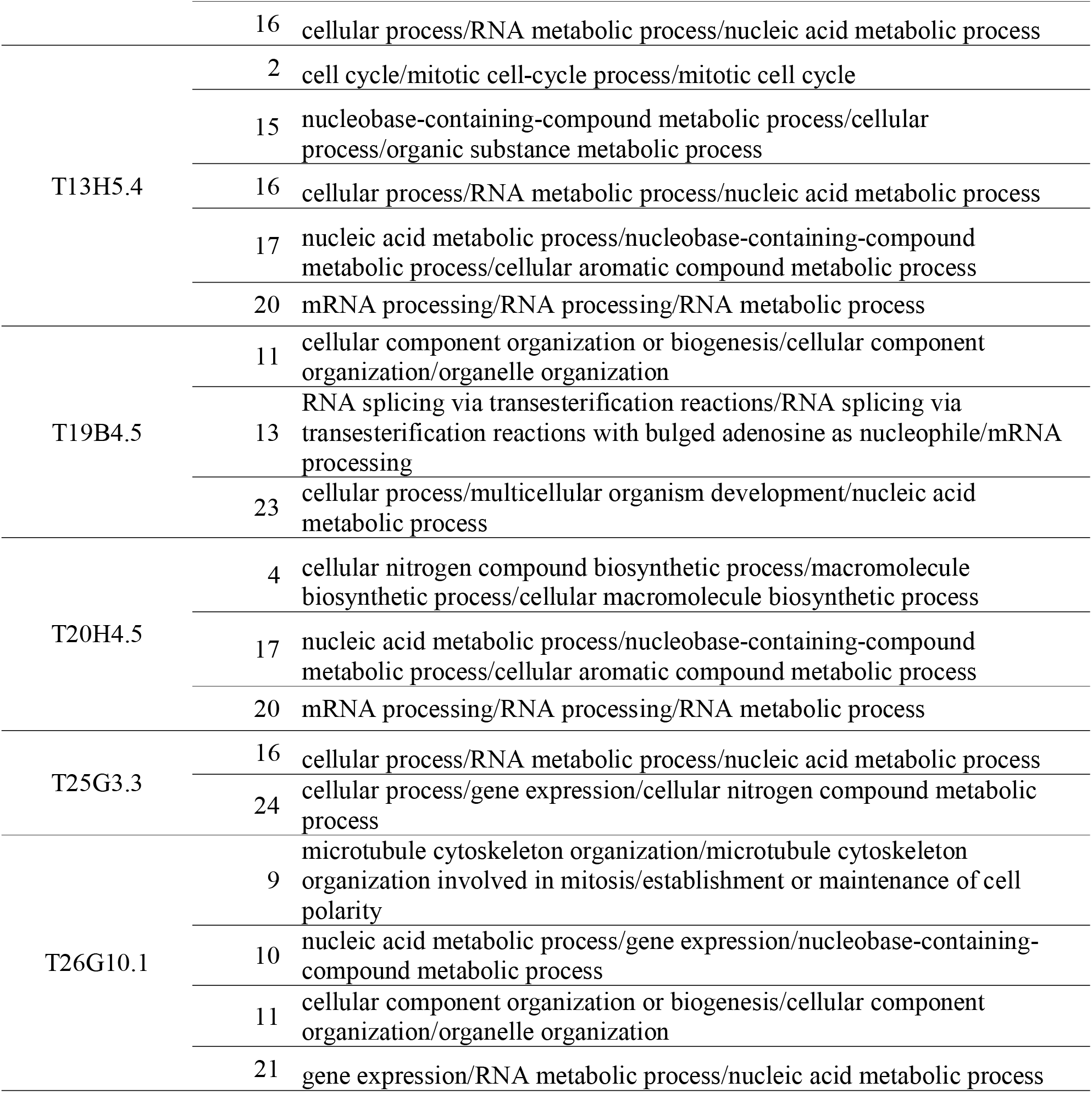
Predicted Biological Process gene ontology terms for the 18 uncharacterized genes

To compare our phenotype-based prediction with the widely used homology-based prediction, we referred to the Gene Ontology database (http://geneontology.org). On the basis of homology-based prediction we found that 72% (13/18) of the uncharacterized genes were annotated with Biological Process GO terms (Table S11); 77% (10/13) of the homology-based predictions were consistent with our phenotype-based predictions, but the homology-based predictions were more detailed than the phenotype-based predictions. Five uncharacterized genes were not annotated on the basis of homology-based prediction. Three of them—C23G10.8, F56C11.5, and T19B4.5—had orthologs only among nematode species, and these orthologs were not well characterized. The remaining two—C02B10.5 and F32H2.6—had a wide range of orthologs among species, including humans. However, C02B10.5 had highly identical orthologs only among nematode species, and these nematode orthologs were not well characterized. F32H2.6 had paralogs that had higher levels of identity to well-characterized homologs than to F32H2.6 itself. Our phenotype-based method thus provides predictions of gene function that are consistent with those made by the homology-based method. The phenotype-based method can predict the functions of genes that cannot be predicted by homology-based methods because of a lack of well-characterized highly identical orthologs.

## DISCUSSION

Here, we created a deep collection of quantitative data on nuclear division dynamics in *C. elegans* embryos RNAi-treated for each of 263 essential embryonic genes. These 263 genes are essential for embryogenesis, and embryos that were RNAi-treated for these genes underwent cell division. We obtained 33 sets of quantitative data from wild-type embryos and five or more sets of quantitative data per gene from RNAi-treated embryos for 189 genes.

A major advantage of our collection is the number of datasets per condition. The availability of five or more datasets per condition enabled us to use a variety of standard statistical analyses. We subjected our collection to a two-sample unequal variance *t*-test followed by multiple test correction and found 8860 RNAi-induced phenotypes for 421 phenotypic characters among the 263 essential embryonic genes. Because the two-sample *t*-test is commonly used in experimental biology, these 8660 RNAi-induced phenotypes will constitute a reliable resource for future experimental analysis.

The method based on data transformation and *P*-value estimation was complementary to that based on a two-sample *t*-test. The former method detected 95% of the previously reported RNAi-induced phenotypes reproduced in our experiment, whereas the latter detected 68% of them. The two methods collectively detected all but one of these phenotypes. The method based on data transformation and *P*-value estimation works well when the penetrance or expressivity of the RNAi silencing is low. It can detect a phenotype even when only one dataset exhibits that phenotype. It can also work well when there are insufficient datasets to apply standard statistical methods such as a two-sample *t-*test. Therefore, the method based on data transformation and *P*-value estimation should be applicable in practical settings to higher organisms such as zebrafish and mice, for which multiple datasets have not been obtained.

Another advantage of our collection is its four-dimensionality: that is, it can capture changes over time and in 3D space. By using our collection, we can objectively detect temporal and 3D spatial phenotypes at a single-cell level. In contrast, it is difficult to objectively detect such phenotypes via manual analysis, because in *C. elegans* embryos most cells divide asynchronously and generate a 3D multicellular structure. Moreover, our collection enables precise comparison of 3D spatial phenotypic characters, because registration of embryos can be performed precisely by using embryonic regions that have been detected from DIC microscopy images. An advantage of DIC microscopy images is that they include not only information about nuclei and spindles but also other information about the embryo (as described below).

Our clustering analysis using 1141 datasets for 263 genes suggests that there are 24 functional processes essential for nuclear division dynamics during the first three rounds of cell division in *C. elegans* embryos. The 161 genes for which fewer than five sets of quantitative data could be obtained (see Figure 1D) exhibited nine RNAi-induced phenotypes: no embryos (sterile); no cell division; and defects in nuclear appearance, spindle appearance, nuclear positioning, nuclear number, cytokinesis, yolk granule density, or embryo size (Table S12). These results suggest that early *C. elegans* embryogenesis can be divided into at least these 33 functional processes, corresponding to the 24 clusters for 262 genes and nine phenotypes for 161 genes.

Many genes were involved in more than one cluster. Many genes have multiple functions (Hodgkin, 1998). Such genes can influence multiple functional processes when they are perturbed. Genes involved in several clusters could act as connecting points between different protein complexes or pathways (Zou et al., 2008).

The top three Biological Process GO terms with the smallest FDR-corrected *P*-values were used to represent each cluster’s functional process. We first tried to annotate each cluster by using the top single GO term. However, the top GO terms for several clusters seemed inappropriate, because the terms were too general and therefore not informative. Although we selected the top three GO terms in this study, it is critical to select an informative subset of GO terms for interpreting the results of the analysis. To achieve a more accurate determination of each cluster’s functional process, we need an index of whether or not each GO term is informative. One possible solution may be to use an automated method of identifying informative subsets of GO terms through word-use profiling of the literature (Jin and Lu, 2010).

The 350 target genes in this study exhibited 100% embryonic lethality in a previous screen (Kamath et al., 2003). However, only 29% of them reproduced this 100% embryonic lethality in our RNAi experiments (Table S1). The difference between our results and the previous results (Kamath et al., 2003) likely originates primarily in the difference in the RNAi method used (injection versus feeding) and in the phenotype scoring methods (e.g., the temperature [22 °C versus 15 °C] and duration [18 h versus 72 h] of RNAi induction) (Piano and Gunsalus, 2002).

In this study, we targeted all essential embryonic genes that were reported to exhibit 100% embryonic lethality in a previous genome-wide screen (Kamath et al., 2003). The previous screen also determined the set of genes that exhibited partial embryonic lethality. Moreover, another other genome-wide RNAi screen found another set of embryonic lethal genes (Sönnichsen et al., 2005). These genes could be the next target in our collection to increase the coverage of genes involved in embryogenesis.

Our WDDD2 database provides not only a collection of quantitative data on nuclear division dynamics but also a collection of 4D DIC microscopy images for all wild-type and RNAi-treated embryos used in this study. Our phenotypic analysis may have detected false positives because of technical issues in the measurement and phenotype detection. However, we can use the microscopy images to check the phenotypes detected from the collection of quantitative data. Moreover, the DIC microscopy images provide information about nuclear and spindle dynamics and cellular membrane and cytoplasmic dynamics. Through the development and use of manual analysis (Schnabel et al., 1997), image-filtering-based methods (Cluet et al., 2014), and machine-learning-based methods (Ounkomol et al., 2018), we expect that information on other dynamics will be obtained.

We expect that many data-driven methods will be developed from our collection, as we have seen in the case of genome sequence data. We also expect that similar large collections of quantitative data will be created for a wide variety of morphological dynamics. Our collection will be a precious resource for understanding the molecular mechanisms of animal development.

## Supporting information

Key Resource Table

## ACKNOWLEDGMENTS

Worm strains were provided by the Caenorhabditis Genetics Center, funded by the NIH National Center for Research Resources. We thank E. Adachi for her contribution to the experiments. We also thank J. Kuramochi, K. Henmi, E. Nagai, K. Shimada, T. Sugimoto, C. Tamai, and S. Yashiro for data curation, and the staff of S.O.’s laboratory for discussions. This work was supported by KAKENHI (Grant-in-Aid for Scientific Research) on Priority Areas ‘Systems Genomics’ [17017038] from the Ministry of Education, Culture, Sports, Science and Technology of Japan (to S.O.); JSPS KAKENHI Grant Number JP18H05412; National Bioscience Database Center, Japan Science and Technology Agency (JST) (to S.O.); and Core Research for Evolutionary Science and Technology Grant Number JPMJCR1511, JST (to S.O).

## AUTHOR CONTRIBUTIONS

K.K. performed the computational analysis and produced the software packages and database. H.O. and K.K. performed the RNAi experiments. H.I. developed the database for sharing microscopy images, quantitative data, and the results of the phenotypic analysis. S.O. conceived and designed the project. K.K. and S.O. wrote the manuscript.

## STAR METHODS

### CONTACT FOR REAGENT AND RESOURCE SHARING

Further information and requests for resources and reagents should be directed to, and will be provided by, the Lead Contact, Shuichi Onami.

### EXPERIMENTAL MODEL AND SUBJECT DETAILS

*Caenorhabditis elegans* nematodes were grown on NGM (Nematode Growth Medium) agar plates at 22 °C in accordance with standard procedures (Brenner, 1974). For all experiments, we used embryos removed from young adult hermaphrodites.

### METHOD DETAILS

#### RNAi

Genes were amplified by PCR from genomic DNA by using the gene-specific primers reported by Kamath et al. (2003). T7 and T3 promoter sequences were added to the 5□Fi-ends of all forward and reverse primers, respectively. By using the PCR product as a template, both RNA strands were simultaneously synthesized with T3 and T7 RNA polymerase (Promega, Madison, WI, USA), and the RNA mixture was heat-denatured and annealed to form dsRNA. Double-strandedness was assessed by scoring shifts in mobility with respect to single-stranded RNA in 0.8% agarose gels. For each gene, dsRNA was injected into between 5 and 24 young adult *C. elegans* (N2 strain) hermaphrodites, as described previously (Mello et al., 1991). The animals were maintained at 22 °C for 18 h in accordance with standard procedures (Brenner, 1974).

#### Embryonic Lethality Test

Five to 10 animals were transferred to individual fresh plates 18 h after dsRNA injection and maintained at 22 °C. After 2 to 4 h, the animals were removed from these plates.

Embryonic lethality was determined by counting all F_1_ progeny at the egg stage (at the time of animal removal) and the larval stage (1 day after animal removal). The number of unhatched eggs was then calculated.

#### Four-dimensional DIC Microscopy Image Recording

Animals removed from the plates in the embryonic lethality test were used for the recordings. A single embryo immediately after fertilization was dissected from each animal and mounted in a well of a multiwell slide (MP Biomedicals, Irvine, CA, USA) coated with 0.01% poly-L-lysine (Sigma-Aldrich, St. Louis, MO, USA) in M9 solution, covered with a coverslip, and sealed with VALAP (Vaseline, lanolin, and paraffin wax melted at equal concentrations). Nomarski DIC microscopy images were obtained with a Leica DM6000 B microscope equipped with an HCX PL APO 63×/1.20 W CORR objective, the illumination intensity and objective-side Wollaston prism of which were adjusted to obtain images of consistent quality. Digital images of 600 × 600 pixels with 256 gray levels (0.1015 μm/pixel) were recorded with a Leica DFC 360 FX CCD camera, and the recording system was controlled by Leica LAS AF6000 software. Digital images of the developing embryo were recorded at 22 °C in 66 focal planes, with 0.5016 μm between focal planes. One set of 66 focal plane images was recorded every 20 s.

### QUANTIFICATION AND STATISTICAL ANALYSIS

#### Measurement of Quantitative Data on Nuclear Division Dynamics

Quantitative data on nuclear division dynamics were obtained from 4D DIC microscopy images of wild-type and RNAi-treated embryos, as described previously (Hamahashi et al., 2007). Errors in detection and tracking of nuclear regions were manually corrected.

#### Dataset Registration and Cell Naming

3D volumetric embryonic regions were detected by using a background subtraction technique and an active contour model for each focal plane. The AP axis was determined by calculating the first principal axis of the 3D volumetric embryonic region. Because the embryo often rotated in the eggshell over time, the DV and LR axes were determined at each time point so that the position of the nucleus of EMS, E, or Ea (the anterior daughter cell of E) became the ventral pole. The embryonic center was determined by calculating the gravity point of the 3D volumetric embryonic region. The use of these embryonic axes and center enabled each dataset to be aligned in a common coordinate system. Cells were named according to the standard nomenclature (Deppe et al., 1978; Sulston et al., 1983).

#### Computational Phenotype Analysis

Four-hundred and twenty-one phenotypic characters were defined mathematically from the pronuclear fusion stage to the eight-cell stage (Table S3). These characters corresponded to the 16 defect categories defined in a previous large-scale manual phenotypic analysis of pronuclear fusion to the four-cell stage (Table S2) (Sönnichsen et al., 2005) but were extended here to the eight-cell stage. The values of these characters were extracted by using an in-house program in the C++ programming language.

To detect RNAi-induced phenotypes, we first used a conventional method based on a two-sample *t*-test. The values of the abovementioned characters were calculated by using the data from wild-type and RNAi-treated embryos, and the *P-*value of each gene for each character was calculated by using Welch’s *t*-test with rank transformation (Ruxton, 2006). For multiple test correction, the q-value of each gene for each character was calculated from the *P*-values by using Storey’s method (Storey and Tibshirani, 2003). RNAi-induced phenotypes were detected by using a threshold q-value of < 0.05.

We next developed a new method to judge whether an RNAi-treated embryo exhibited an alteration of phenotypic character, by estimating *P*-values on the basis of the distribution of values of phenotypic characters in wild-type embryos, as follows. The values of the phenotypic characters in wild-type embryos were transformed by using the Johnson transformation method (Chou et al., 1998). The value of the character in each RNAi-treated embryo was then transformed by using the parameters of Johnson transformation determined in the data from wild-type embryos. The transformed data for each of the phenotypic characters followed a normal distribution (Shapiro-Wilk test *P*-value > 0.05). The threshold *P*-value for alteration of phenotypic character was set to 0.001 because the use of this threshold did not detect any alteration in phenotypic data in wild-type embryos. These two statistical analyses were performed by using an inhouse program in R statistical language (R Core Team, 2014).

#### Clustering Analysis

To overcome the different scale and units of the 421 phenotypic characters, the value of each phenotypic character extracted from an RNAi-treated embryo was normalized as follows: *v*=(*x* – mean_wt_) / sd_wt_, where *v* is the normalized value, *x* is the original value extracted from the RNAi dataset, and mean_wt_ and sd_wt_ are, respectively, the mean value and standard deviation of the values extracted from wild-type embryos. After the normalization, one of the *lam-2(RNAi*) embryos produced outliers for the ratio of the volume of EMS to P_1_ and the volume of P_2_ to P_1_; this embryo was omitted from further analysis. Therefore, 1141 out of the 1142 RNAi-treated embryos were used for the cluster analysis. Next, the missing values of phenotypic characters were estimated by using the KNNimpute method (Troyanskaya et al., 2001). Imputation was performed with the impute package (https://www.bioconductor.org/packages/release/bioc/html/impute.html) in R statistical language (R Core Team, 2014). After the above preprocessing, hierarchical clustering by using the Euclidean metric and Ward’s linkage was applied to the phenotypic expression profiles of the 1141 RNAi-treated embryos. The clustering analysis was performed with an in-house program in Python.

#### GO Enrichment Analysis

To annotate each gene cluster with specific biological processes, GO enrichment analysis was performed. We first selected genes for which multiple RNAi-treated embryos were grouped in a cluster. We next found the top three Biological Process GO terms that were overrepresented in the genes in the cluster by using the GO enrichment analysis tool (FDR < 0.05) provided by the Gene Ontology Consortium (Mi et al., 2013).

### DATA AND SOFTWARE AVAILABILITY

All image and quantitative data are available through the Worm Developmental Dynamics Database 2 (WDDD2; https://wddd.riken.jp). The results of the phenotypic analysis and clustering analysis are also available at the WDDD2. All in-house programs for data analysis are available at https://github.com/wddd-tools/phenotype_analysis/.

---

**Figure S1.** Features of RNAi-induced phenotypes in early *Caenorhabditis elegans* embryogenesis, as detected by using a method based on data transformation and *P*-value estimation.

**Table S1.** Embryonic lethality in this study.

**Table S2.** Defined phenotypic characters corresponding to the 16 defect categories in the previous manual analysis.

**Table S3.** The 421 phenotype characters defined in this study.

**Table S4.** Phenotypic expression profiles of wild-type and RNAi-treated embryos.

**Table S5.** Results of computational analysis by using Welch’s *t*-test with rank transformation.

**Table S6.** Results of computational analysis by using data transformation and *P*-value estimation.

**Table S7.** Results of gene ontology (GO) analysis for genes counted in the right narrow peak and in the vicinity of the right narrow peak.

**Table S8.** Genes in the 24 clusters.

**Table S9.** Top three Biological Process gene ontology GO terms for each of the 24 gene clusters.

**Table S10.** Genes and their descriptions in Cluster 9 and Cluster 8.

**Table S11.** Biological Process gene ontology (GO) terms predicted for the 18 uncharacterized genes.

**Table S12.** RNAi-induced phenotypes for the 161 genes for which fewer than five sets of quantitative data could be obtained.

## REFERENCES

Andralojc, K. M., Campbell, A. C., Kelly, A. L., Terrey, M., Tanner, P. C., Gans, I. M., Senter-Zapata, M. J., Khokhar, E. S., and Updike, D. L. (2017). ELLI-1, a novel germline protein, modulates RNAi activity and P-granule accumulation in *Caenorhabditis elegans*. PLoS Genet. 13:e1006611.

Bao, Z., Murray, J.I., Boyle, T., Ooi, S.L., Sandel, M.J., and Waterston, R.H. (2006). Automated cell lineage tracing in *Caenorhabditis elegans*. Proc. Natl. Acad. Sci. USA 103, 2707–2712.

Bashar, M.K., Komatsu, K., Fujimori, T., and Kobayashi, T.J. (2012). Automatic extraction of nuclei centroids of mouse embryonic cells from fluorescence microscopy images. PLoS One 7, e355550.

Brenner, S. (1974). The genetics of *Caenorhabditis elegans*. Genetics 77, 71–94.

Camon, E.B., Barrell, D.G., Dimmer, E.C., Lee, V., Magrane, M., Maslen, J., Binns, D., and Apweiler, R. (2005). An evaluation of GO annotation retrieval for BioCreAtIvE and GOA. BMC Bioinformatics 6 Suppl 1:S17.

Chou, Y.M., Polansky, A.M., and Mason, R.L. (1998). Transforming non-normal data to normality in statistical process control. J. Qual. Technol. 30, 133–141.

Claycomb, J. M., Batista, P. J., Pang, K. M., Gu, W., Vasale, J. J., van Wolfswinkel, J. C., Chaves, D. A., Shirayama, M., Mitani, S., Ketting, R. F., Conte, D. Jr., and Mello, C. C. (2009). The Argonaute CSR-1 and its 22G-RNA cofactors are required for holocentric chromosome segregation. Cell 139, 123–134.

Cluet, D., Stébé, P.N., Riche, S., Spichty, M., and Delattre, M. (2014). Automated high-throughput quantification of mitotic spindle positioning from DIC movies of Caenorhabditis embryos. PLoS One 9, e93718.

Deppe, U., Schierenberg, E., Cole, T., Krieg, C., Schmitt, D., Yoder, B., and von Ehrenstein, G. (1978). Cell lineage of the embryo of the nematode *Caenorhabditis elegans*. Proc. Natl. Acad. Sci. USA 75, 376–380.

Dittman, J. S., and Kaplan, J. M. (2008). Behavioral impact of neurotransmitter-activated G-protein-coupled receptors: muscarinic and GABAB receptors regulate Caenorhabditis elegans locomotion. J. Neurosci. 28, 7104–7112.

Fire, A., Xu, S., Montgomery, M.K., Kostas, S.A., Driver, S.E., and Mello, C.C. (1998) Potent and specific interference by double-stranded RNA in *Caenorhabditis elegans*. Nature 391, 806–811.

George-Raizen, J.B., Shockley, K.R., Trojanowski, N.F., Lamb, A.L., and Raizen, D.M. (2014). Dynamically-expressed prion-like proteins form a cuticle in the pharynx of *Caenorhabditis elegans*. Biol. Open 3, 1139–1149.

Hamahashi, S., Onami, S., and Kitano, H. (2005). Detection of nuclei in 4D Nomarski DIC microscope images of early *Caenorhabditis elegans* embryos using local image entropy and object tracking. BMC Bioinformatics 6:125.

Hamahashi, S., Kitano, H., and Onami, S. (2007). A system for measuring cell division patterns of early *Caenorhabditis elegans* embryos by using image processing and object tracking. Systems Comput. Jpn. 38, 12–24.

Ho, V.W., Wong, M.K., An, X., Guan, D., Shao, J., Ng, H.C., Ren, X., He, K., Liao, J., Ang, Y., Chen, L., Huang, X., Yan, B., Xia, Y., Chan, L.L., Chow, K.L., Yan, H., and Zhao, Z. (2015). Systems-level quantification of division timing reveals a common genetic architecture controlling asynchrony and fate asymmetry. Mol. Syst. Biol. 11, 814.

Hodgkin, J. (1998). Seven types of pleiotropy. Int. J. Dev. Biol. 42, 501–505.

Hyun, M., Bohr, V.A., and Ahn, B. (2008). Biochemical characterization of the WRN-1 RecQ helicase of Caenorhabditis elegans. Biochemistry 47, 7583–7593.

Jin, B., and Lu, X. (2010) Identifying informative subsets of the Gene Ontology with information bottleneck methods. Bioinformatics 26, 2445–2451.

Kamath, R.S., Fraser, A.G., Dong, Y., Poulin, G., Durbin, R., Gotta, M., Kanapin, A., Le Bot, N., Moreno, S., Sohrmann, M., Welchman, D.P., Zipperlen, P., and Ahringer, J. (2003). Systematic functional analysis of the *Caenorhabditis elegans* genome using RNAi. Nature 421, 231–237.

Keller, P.J., Schmidt, A.D., Wittbrodt, J., and Stelzer, E.H. (2008). Reconstruction of zebrafish early embryonic development by scanned light sheet microscopy. Science 322, 1065–1069.

Keller, P.J., Schmidt, A.D., Santella, A., Khairy, K., Bao, Z., Wittbrodt, J., and Stelzer, E.H. (2010). Fast, high-contrast imaging of animal development with scanned light sheet-based structured-illumination microscopy. Nat. Methods 7, 637–642.

Kim, W., Underwood, R.S., Greenwald, I., and Shaye, D.D. (2018). OrthoList 2: a new comparative genomic analysis of human and *Caenorhabditis elegans* genes. Genetics 210, 445–461.

Kyoda, K., Adachi, E., Masuda, E., Nagai, Y., Suzuki, Y., Oguro, T., Urai, M., Furukawa, M., Shimada, K., Kuramochi, J., Nagai, E., and Onami, S. (2013). WDDD: Worm Developmental Dynamics Database. Nucleic Acids Res. 41(Database issue), D732–D737.

Leidel, S., Delattre, M., Cerutti, L., Baumer, K., and Gonczy, P. (2005). SAS-6 defines a protein family required for centrosome duplication in *C. elegans* and in human cells. Nat. Cell Biol. 7, 115–125.

Lu, X. and Horvitz, H.R. (1998) lin-35 and lin-53, two genes that antagonize a *C. elegans* Ras pathway, encode proteins similar to Rb and its binding protein RbAp48. Cell 95, 981–991.

Mello, C.C., Kramer, J.M., Stinchcomb, D., and Ambros, V. (1991). Efficient gene transfer in *C. elegans:* extrachromosomal maintenance and integration of transforming sequences. EMBO J 10, 3959–3970.

Mi, H., Muruganujan, A., Gasagrande, J.T., and Thomas, P.D. (2013). Large-scale gene function analysis with the PANTHER classification system. Nat. Protoc. 8, 1551–1566.

Mitchell, P. (1961). Coupling of phosphorylation to electron and hydrogen transfer by a chemi-osmotic type of mechanism. Nature 191, 144–148.

Neumann, B., Walter, T., Hériché, J. K., Bulkescher, J., Erfle, H., Conrad, C., Rogers, P., Poser, I., Held, M., Liebel, U., Cetin, C., Sieckmann, F., Pau, G., Kabbe, R., Wünsche, A., Satagopam, V., Schmitz, M. H., Chapuis, C., Gerlich, D. W., Schneider, R., Eils, R., Huber, W., Peters, J.M., Hyman, A.A., Durbin, R., Pepperkok, R., and Ellenberg, J. (2010). Phenotypic profiling of the human genome by time-lapse microscopy reveals cell division genes. Nature 464, 721–727.

Ohya, Y., Sese, J., Yukawa, M., Sano, F., Nakatani, Y., Saito, T.L., Saka, A., Fukuda, T., Ishihara, S., Oka, S., Suzuki, G., Watanabe, M., Hirata, A., Ohtani, M., Sawai, H., Fraysse, N., Latgé, J.P., François, J.M., Aebi, M., Tanaka, S., Muramatsu, S., Araki, H., Sonoike, K., Nogami, S., and Morishita, S. (2005). High-dimensional and large-scale phenotyping of yeast mutants. Proc. Natl. Acad. Sci. USA 102, 19015–19020.

Ounkomol, C., Seshamani, S., Maleckar, M.M., Collman, F., and Johnson, G.R. (2018). Label-free prediction of three-dimensional fluorescence images from transmitted-light microscopy. Nat. Methods 15, 917–920.

Piano, F. and Gunsalus, K.C. (2002). RNAi-based functional genomics in *Caenorhabditis elegans*. Curr. Genomics 3, 68–81.

Poulin, G., Dong, Y., Fraser, A.G., Hopper, N.A., Ahringer, J. (2005). Chromatin regulation and sumoylation in the inhibition of Ras-induced vulval development in *Caenorhabditis elegans*. EMBO J 24, 2613–2623.

R Core Team. (2014). R: a language and environment for statistical computing. R Foundation for Statistical Computing, Vienna, Austria. URL http://www.R-project.org/.

Rappleye, C.A., Tagawa, A., Lyczak, R., Bowerman, B, and Aroian, R.V. (2002). The anaphase-promoting complex and separin are required for embryonic anterior-posterior axis formation. Dev. Cell 2, 195–206.

Ruxton, G.D. (2006). The unequal variance *t*-test is an underused alternative to Student’s *t*-test and the Mann-Whitney *U* test. Behav. Ecol. 17, 688–690.

Santella, A., Kovacevic, I., Herndon, L.A., Hall, D.H., Du, Z., and Bao, Z. (2016). Digital development: a database of cell lineage differentiation in *C. elegans* with lineage phenotypes, cell-specific gene fucntions and a multiscale model. Nucleic Acids Res. 44(D1), D781–D785.

Schnabel, R., Hutter, H., Moerman, D., Schnabel, H. (1997). Assessing normal embryogenesis in *Caenorhabditis elegans* using a 4D microscope: variability of development and regional specification. Dev. Biol. 184, 234–265.

Seydoux, G. (2018). The P Granules of *C. elegans:* A Genetic Model for the Study of RNA-Protein Condensates. J. Mol. Biol. 430, 4702–4710.

Sönnichsen, B., Koski, L.B., Walsh, A., Marschall, P., Neumann, B., Brehm, M., Alleaume, A.M., Artelt, J., Bettencourt, P., Cassin, E., Hewitson, M., Holz, C., Khan, M., Lazik, S., Martin, C., Nitzsche, B., Ruer, M., Stamford, J., Winzi, M., Heinkel. R., Roder, M., Finell, J., Hantsch, H., Jones, S.J., Jones, M., Piano, F., Gusalus, K.C., Oegema, K., Gonczy, P., Coulson, A., Hymann, A.A., and Echeverri, C.J. (2005). Full-genome RNAi profiling of early embryogenesis in *Caenorhabditis elegans*. Nature 434, 462–469.

Storey, J.D. and Tibshirani, R. (2003). Statistical significance for genomewide studies. Proc. Natl. Acad. Sci. USA 100, 9440–9445.

Sulston, J.E., Schierenberg, E., White, J.G., and Thomson, J.N. (1983). The embryonic cell lineage of the nematode *Caenorhabditis elegans*. Dev. Biol. 100, 64–119.

Tabuse, Y., Izumi, Y., Piano, F., Kemphues, K.J., Miwa, J., and Ohno, S. (1998). Atypical protein kinase C cooperates with PAR-3 to establish embryonic polarity in *Caenorhabditis elegans*. Development 125, 3607–3614.

Troyanskaya, O., Cantor, M., Sherlock, G., Brown, P., Hastie, T., Tibshirani, R., Botstein, D., and Altman, R.B. (2001) Missing value estimation methods for DNA microarrays. Bioinformatics 17, 520–525.

Yoneda, T., Benedetti, C., Urano, F., Clark, S.G., Harding, H.P., and Ron, D. (2004). Compartment-specific perturbation of protein handling activates genes encoding mitochondrial chaperones. J. Cell Sci. 117, 4055–4066.

Zou, L., Sriswasdi, R., Ross, B., Missiuro, P.V., Liu, J., and Ge, H. (2008). Systematic analysis of pleiotropy in *C. elegans* early embryogenesis. PLoS Comput. Biol., 4, e1000003.

