## Supplementary material for "Deep Collection of Quantitative Nuclear Division Dynamics Data in RNAi-treated *Caenorhabditis elegans* Embryos": Key Resource Table

**KEY RESOURCES TABLE**

| REAGENT or RESOURCE | SOURCE | IDENTIFIER |
| --- | --- | --- |
| *Chemicals, Peptides, and Recombinant Proteins* | | |
| T3, T7 RNA polymerase | Promega | P208C, P207B |
| poly-L-lysine | Sigma-Aldrich | P8920–100ML |
| *Experimental Models: Organism/Strain* | | |
| *Caenorhabditis elegans*: N2 | CGC (Caenorhabditis Genetics Center) | <https://cgc.umn.edu/strain/N2> |
| *Oligonucleotides* | | |
| PCR primers | SJJ set in Kamath et al., 2003 | N/A |
| *Deposited Data* | | |
| Raw image data | This paper | [https://wddd.riken.jp](https://wddd.riken.jp/) |
| Quantitative data  (processed data) | This paper | [https://wddd.riken.jp](https://wddd.riken.jp/) |
| Embryonic lethality test | This paper | [https://wddd.riken.jp](https://wddd.riken.jp/) |
| Phenotypic analysis data | This paper | [https://wddd.riken.jp](https://wddd.riken.jp/) |
| *Software and Algorithms* | | |
| Image-processing tool | Hamahashi et al., 2007 | N/A |
| Phenotypic analysis tools | This paper | <https://github.com/wddd-tools/phenotype_analysis>/ |
| jtrans | CRAN | <https://cran.r-project.org/web/packages/jtrans/> |
| impute | Bioconductor | <https://www.bioconductor.org/packages/release/bioc/html/impute.html> |
| Clustering tool | This paper | <https://github.com/wddd-tools/phenotype_analysis/> |
