## Supplementary material for "Deep Collection of Quantitative Nuclear Division Dynamics Data in RNAi-treated *Caenorhabditis elegans* Embryos": Figure S1

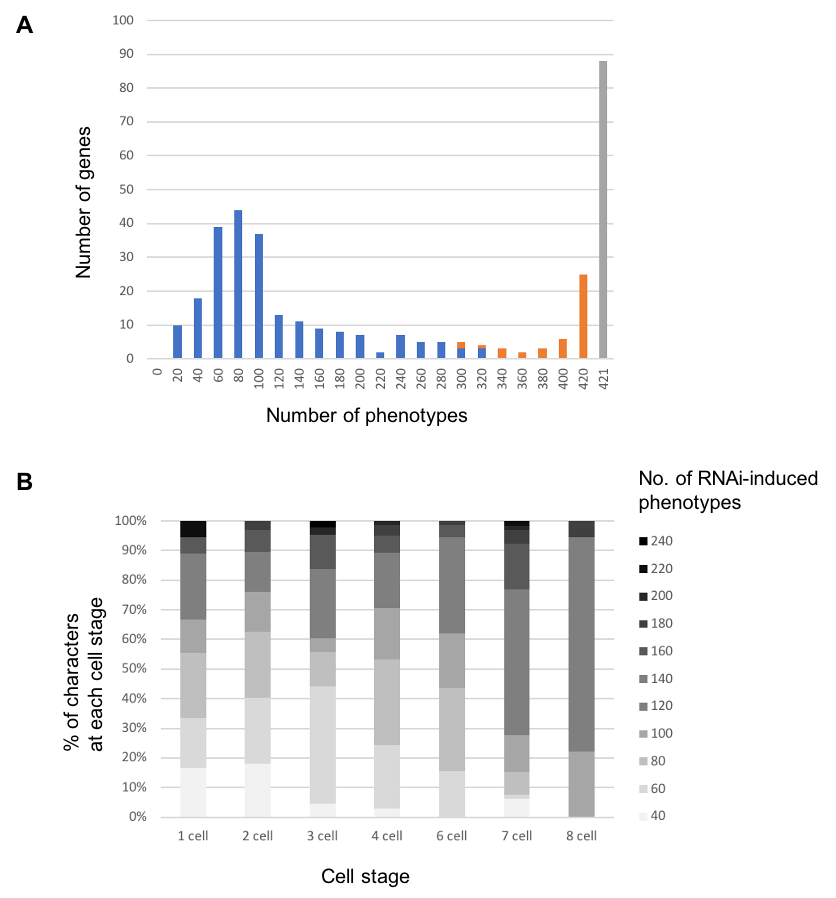

**Figure S1.** Features of RNAi-induced phenotypes in early *Caenorhabditis elegans* embryogenesis, as detected by using a method based on data transformation and *P*-value estimation. Overall, the results showed features similar to those of the results of the two-sample *t*-test-based method (see Fig. 3C). (A) Histogram of the number of genes with the indicated numbers of RNAi-induced phenotypes among the 421 phenotypic characters. Bins 0, 20, … , 420, 421 represent 0, 1 to 20, … , 401 to 420, 421, respectively. Blue, orange, and gray represent genes for which we could calculate more than half, fewer than half, and none, respectively, of the 421 phenotypic characters in the RNAi-treated embryos. (B) Distribution of the number of RNAi-induced phenotypes for each phenotypic character at each cell stage (one-cell to eight-cell).
